# Hypertension drives microbial translocation and shifts in the fecal microbiome of non-human primates

**DOI:** 10.1101/2021.07.30.454379

**Authors:** Ravichandra Vemuri, Alistaire Ruggiero, Jordyn B. Whitfield, Greg O. Dugan, J. Mark Cline, Masha R. Block, Hao Guo, Kylie Kavanagh

**Affiliations:** Department of Pathology, Section on Comparative Medicine, Wake Forest School of Medicine, Medical Center Boulevard, Winston-Salem, NC 27157, USA; Lineberger Comprehensive Cancer Center, University of North Carolina at Chapel Hill, Chapel Hill, NC, USA; Department of Genetics, University of North Carolina at Chapel Hill, Chapel Hill NC, USA; Department of Biomedicine, University of Tasmania, Hobart, 7000, TAS, Australia

**Keywords:** blood pressure, hypertension, microbial translocation, gut barrier dysfunction, dysbiosis

## Abstract

Accumulating evidence indicates a link between gut barrier dysfunction and hypertension. However, it is unclear whether hypertension dictates gut barrier dysfunction or vice versa and whether the gut microbiome plays a role. To better understand this relationship, first, we cross-sectionally examined hypertension and other cardio-metabolic risk factors and gut barrier function in a population of 150 nonhuman primates. Interestingly, the animals with hypertension showed evidence of gut barrier dysfunction (i.e., translocation of microbes through the gut wall), as indicated by higher plasma levels of lipopolysaccharide-binding protein (LBP)-1, compared to normotensive animals. Further, plasma LBP-1 levels were strongly correlated with diastolic blood pressure, independent of age and other health markers, suggesting specificity of the effect of hypertension on microbial translocation. In a subsequent longitudinal study (analysis at baseline, 12 and 27 months), hypertensive animals had higher plasma levels of LBP-1 at all the time points and greater bacterial gene expression in mesenteric lymph nodes compared to normotensive animals, confirming microbe translocation. Concomitantly, we identified distinct dysbiosis in the gut microbial signature of hypertensive versus normotensive animals at 12 and 27 months. These results suggest that hypertension drives microbial translocation in the gut and eventually unhealthy shifts in the gut microbiome, possibly contributing to poor health outcomes, providing further impetus for the management of hypertension.

**Graphical Abstract:** 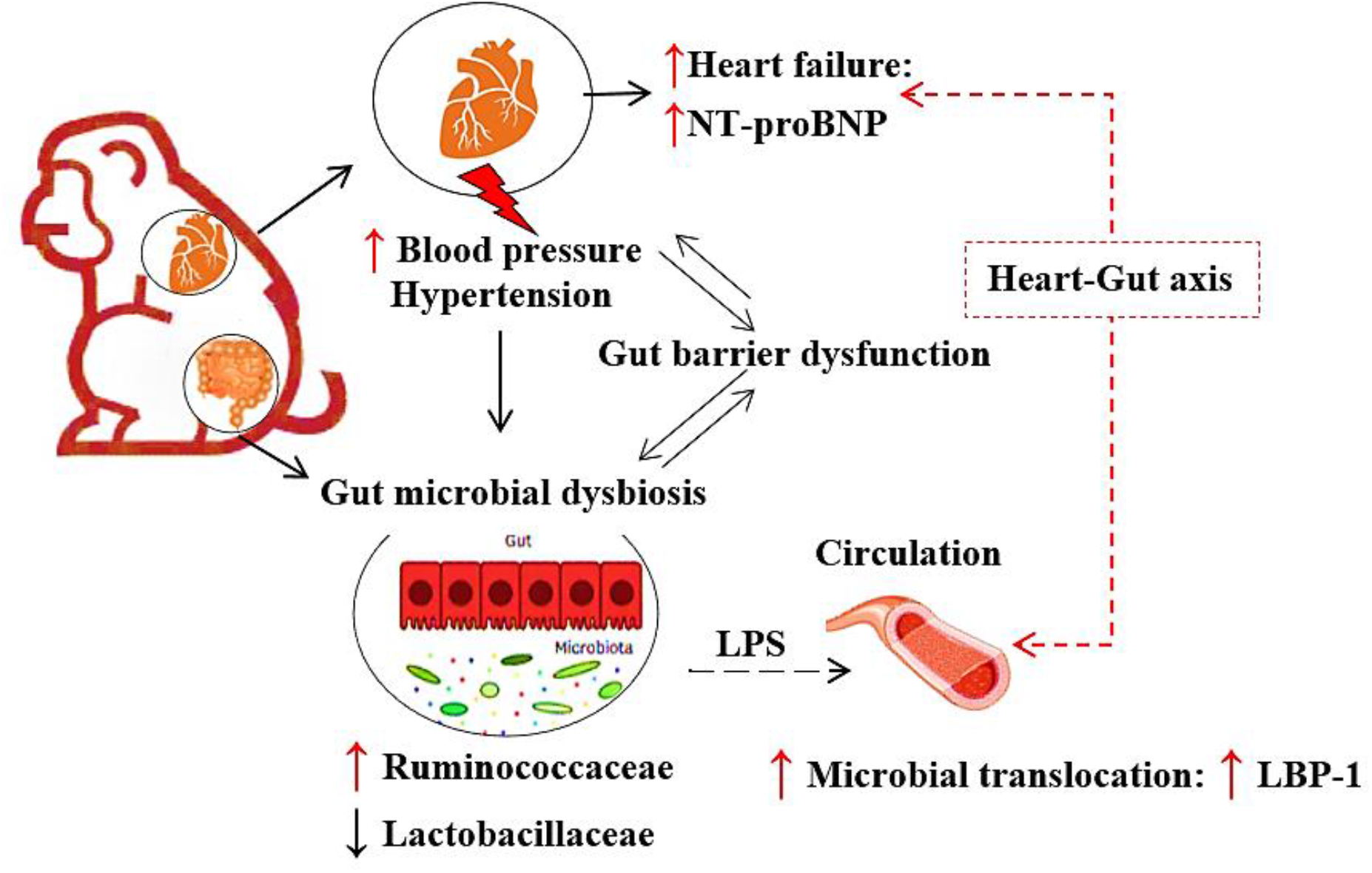

## Introduction

Gut barrier dysfunction and related diseases cost the US healthcare system over $100 billion per year, yet the causes are mostly unknown.^1, 2^ A compromised gut barrier makes the gut mucosal lining “leaky”, enabling the translocation of microbes or microbial products into the systemic circulation from the gut lumen in a process known as microbial translocation (MT), leading to systemic low-grade endotoxemia and inflammation.^3–5^ Further, studies have shown that a defective gut barrier alters the composition of the microbes residing in the gut (gut microbiome), resulting in dysbiosis, which worsens overall health.^1, 3, 6–9^ Therefore, it is essential to comprehend regulators of the gut barrier and gut microbiome-related changes to help prevent the onset of various chronic diseases.

One possible contributor to MT and dysbiosis is risk factors for cardiovascular disease, notably hypertension.^6, 10, 11^ Recently, a study on the spontaneous hypertensive rat model showed increased gut permeability and inflammation, suggesting a possible link between hypertension and gut barrier function.^12, 13^ However, it is unclear whether hypertension dictates gut barrier dysfunction and whether the gut microbiome plays a role. Given the complex nature of these associations, larger cross-sectional studies and longitudinal studies are required to understand the nature of the associations. A further limitation of the existing studies in rats is that they only looked at the gut microbiome at one time point, but this is inadequate to assess the gut microbiome, because other research shows the gut microbiome can change dramatically from one time point to another. Changes in the microbiome result from factors such as aging, environment, diet, activity and other influences, and so looking at the gut microbiome at a number of time points is imperative to capture the true effects.^4, 5, 9^ Due to rapid shifts in the gut microbiome composition and to assess whether the gut barrier and gut microbiota are linked to hypertension, more longitudinal studies are needed.

Animal models are valuable as they can be controlled and used in ways human studies cannot, in terms of age, diet and other environmental factors.^14^ While the rodent models are indisputably useful, they may not recapitulate the human conditions to the same degree as nonhuman primate (NHP) models.^14, 15^ In addition, unlike rodents, NHPs share >93% similarities with the human genome and faithfully recapitulate eating, sleeping and aging patterns.^4, 5, 15, 16^ For this purpose, we selected a NHP model, which spontaneously display several key features of human pathophysiology, including hypertension.^14^ Therefore, in the present study, we aimed to: (i) cross-sectionally investigate the relationship between MT and hypertension (study 1), and (ii) longitudinally examine markers of health, hypertension, gut barrier function, and microbiome in a controlled cohort of NHPs (study 2).

## Material and Methods

### Animals, diet, and study design

For the cross-sectional study (study 1; left panel of Figure 1), we utilized vervet monkeys (*Chlorocebus aethiops sabaeus*) of both sexes, residing in a breeding colony at Wake Forest School of Medicine, NC USA. We used the definition of hypertension based on the American Heart Association’s blood pressure measures, systolic blood pressure (SBP) and diastolic blood pressure (DBP) threshold of 120/80 mmHg^17^ and similarly confirmed in NHPs.^18^ For the longitudinal study (study 2; right panel of Figure 1, rhesus monkeys (*Macaca mulatta*) were used at Wake Forest School of Medicine. These animals were divided as hypertensive and normotensives based on repeated assessments of blood pressure measures and grouped based on the definitions given above. Animals were housed either singly or socially in standard primate enclosures and were maintained at 21–26°C with ~50%−65% relative humidity with 12-hour light and dark periods. Ten NHPs in this longitudinal study were exposed to 4 Gy total body irradiation (TBI) as previously described^19^ which had no effects on study outcomes (Table S3 and Figure S4). Animals were fed a commercial laboratory primate chow (Laboratory Diet 5038 study 1 or 5LOP study 2; LabDiet, St. Louis, MO) with daily supplemental fresh fruits and vegetables. All procedures were performed in accordance with the Guide for Care and Use of Laboratory Animals (IACUC #A18-185 and #A18-062) and in compliance with the Institutional Animal Care and Use Committee, at our Association for the Assessment and Accreditation of Laboratory Animal Care International accredited institution, which operates in compliance with the Animal Welfare Act.

**Figure 1:**
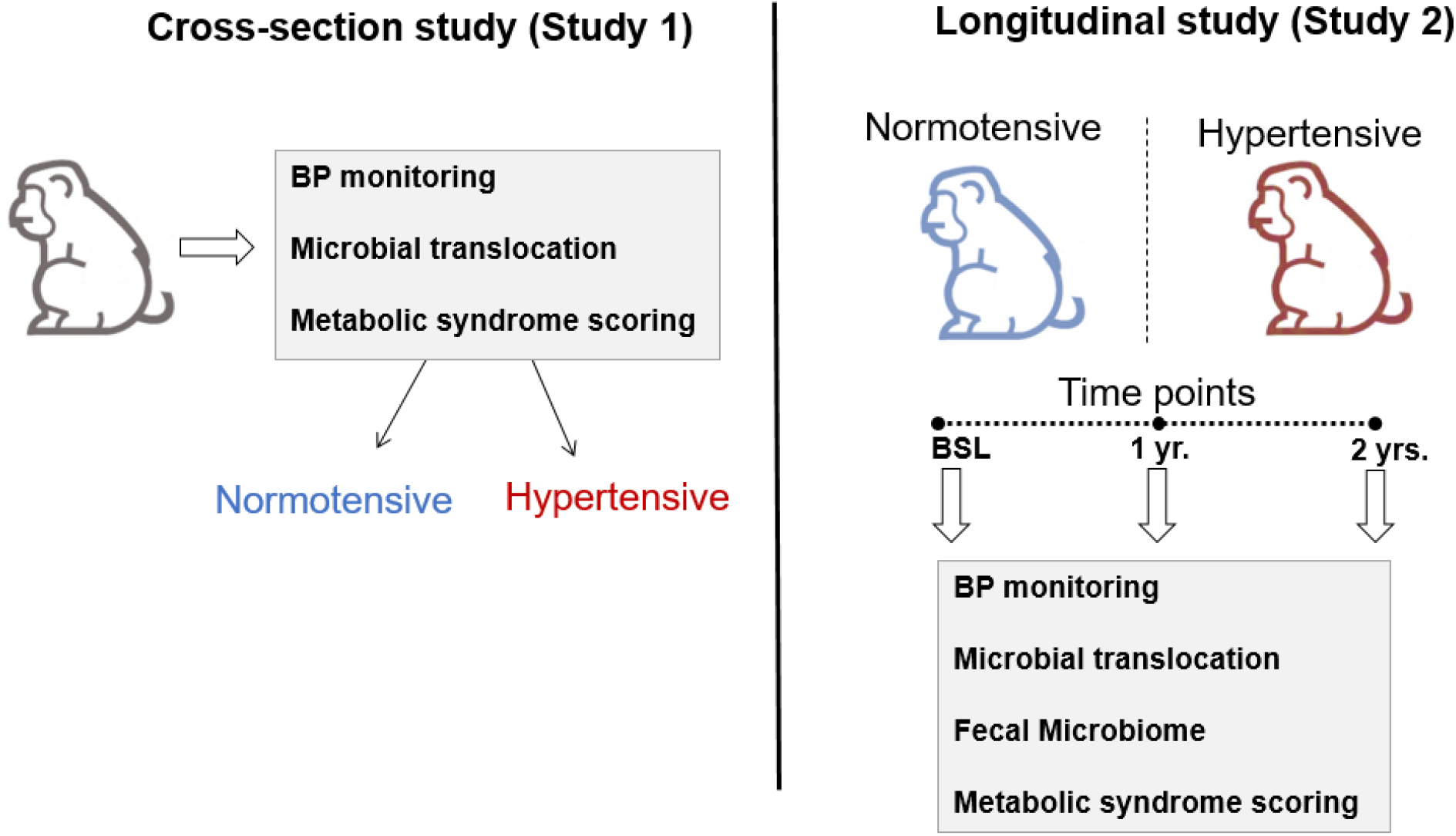
Overall study design. This schematic represents the overview of the 2 studies described in the methods section. BP= blood pressure, BSL= baseline.

### Blood pressure measurement

The blood pressure was measured indirectly using a sphygmomanometer at 15–30 and 40–50 min after ketamine anesthesia (10 mg/kg, i.m.) and was calculated as the mean of 3 measurements.^18^

### Circulating biomarkers and metabolic health markers

Animals were fasted overnight and anesthetized with intramuscular ketamine (10 to 15 mg/kg) to allow for sample and data collections for study 1, and for study 2 samples were collected at baseline, 12 months and 27 months. Each animal was weighed and waist circumference measured with a flexible tape measure at the level of the umbilicus. Blood samples were obtained by venipuncture of the femoral vein into ethylenediaminetetraacetic acid (EDTA) blood tubes. The EDTA anticoagulated blood was held on ice until it could be processed. After processing the plasma, samples were stored at −80 °C until analysis. General metabolic health and biochemistry panels including total plasma triglyceride (TG), high-density lipoprotein cholesterol (HDLC) and total plasma cholesterol (TPC) concentrations, were measured as previously described.^5^ Plasma was used to measure fasting blood glucoses, MT biomarkers, lipopolysaccharide-binding protein (LBP)-1 (Hycult Biotech Inc., Plymouth Meeting, PA) and soluble (s) CD14, and cardia biomarker NT-proBNP (N-terminal pro-B-type natriuretic peptide; MyBiosource, San Diego, CA) via ELISA and following manufacturer instructions as previously described.^4, 5^

### Echocardiogram

Echocardiography was performed on each of the vervet monkeys in study 1 using a GE Logiq S8 ultrasound unit (GE Healthcare, Chicago, IL) using an approach that has been previously described.^20^ Cardiac morphometric and functional phenotypes include: left ventricular diametric at end diastole (LVDd), left ventricular diameter at end systole (LVDs), left ventricular wall fractioning shortening percentage (LVDFS%) calculated as (LVDd-LVDs)/LVDd) x 100 and ejection fraction percentage (EF%). The animal’s body weight (kg) was obtained at the time of echocardiography and used to calculate body surface area (BSA, m^2^) as BW^2/3^ x 0.0969. Echocardiographic analysis was performed using motion (M) mode, parasternal long axis, tissue doppler, 2-chamber, 4-chamber, and 5-chamber views. TOMTEC software (TOMETEC, Chicago, IL) was used for echocardiographic analysis of motion (M) mode, parasternal long axis, tissue doppler, 2-chamber, 4-chamber, and 5-chamber views.

### Fecal sampling and 16S rRNA gene sequencing and data analysis

Fecal samples from the 16 monkeys included in the longitudinal study (study 2) were collected for microbiome characterization and the fecal DNA extracted as described previously.^4, 5, 21^ Briefly, DNA was extracted from fecal samples (Qiagen, Germantown, MD) and the quality of DNA determined by gel electrophoresis. Fecal DNA samples were amplified by PCR using barcoded primer pairs targeting the V3-V4 region of the bacterial 16S gene. PCR amplicons were sequenced using the Mi-Seq Illumina sequencer. The resulting bacterial sequence fragments were clustered into Operational Taxonomic Units (OTUs) and aligned to Greengenes microbial gene database with 97% sequence similarity in QIIME (1.8.0). Bacterial taxonomy summarization, rarefaction analyses of microbial diversity, compositional differences (dissimilarity value indicated by Unweighted UniFrac Distance) were calculated in QIIME as previously described using scripts (including pick_open_reference_otus.py, summarize_taxa.py, alpha_rarefaction.py, jackknifed_beta_diversity.py and make_distance_boxplots.py). PCoA plots were generated by QIIME script (make_2d_plots.py) and each point represents one animal (Figure S4), represent the interquartile range (IQR) during the rarefaction analyses (see http://www.wernerlab.org/teaching/qiime/overview for scripts details).

### Statistical analysis

Statistical analyses were performed using Graphpad Prism 10 (San Diego, CA) and Statistica version 13 (StatSoft, Tulsa, OK). Data are presented as means ± standard error of the mean (SEM) for each group and p values of < 0.05 were considered significant. Metabolic and phenotypic variables were analyzed for differences by analysis of covariance using baseline values, if available, and current body weight. Repeated measures of ANOVA was used for values measured during the 2 years of longitudinal study (study 2). Fisher’s exact test was used to measure differences between proportions, and Pearson’s correlation coefficients were calculated to measure strength of the associations. Multiple regression modeling with backwards stepwise removal of variables was used to find predictors of MT in study 1. TBI exposure was included as a model factor for study 2 analyses, but was not retained due to lack of significant effects.

## Results

### Study 1: Cross-sectional study to understand the relationship between hypertension and MT

Detailed information on demographic and metabolic health biomarkers is shown in Table S1. A total of 153 vervet monkeys, with 15.6 % males (N=24) and 84.3% females (N=129) were evaluated in this study. We observed no differences in any of the measured parameters based on sex. The mean SBP (Figure 2A) and DBP (Figure 2B) of the hypertensive animals (143 ± 2.11 and 83 ± 2.11, respectively) were significantly higher than those of the normotensive animals (102 ± 1.26 and 60 ± 1.16, respectively) (p<0.0001). To understand the role of hypertension on the gut barrier and cardiac failure risk, we measured MT via LBP-1 and sCD14, and cardiac biomarker via NT-proBNP in plasma samples. Hypertensive animals had numerically but non-significantly higher levels of LBP-1 (p=0.06) (Figure S1) compared to the normotensive animals. We observed significantly higher plasma concentrations of NT-proBNP in hypertensive animals (Figure 2C), which is an indicator of strain on cardiomyocytes and of high interest as a prognosticator of heart failure. Consistently, echocardiogram data showed significantly higher E/A (Figure 2D) and % FS (Figure 2E) compared to normotensive animals. This data is an important phenotype as it relates to diastolic dysfunction and risk of heart failure particularly that described as heart failure with preserved ejection fraction. These changes in hypertensive animals showed dramatic remodeling, and almost no left ventricular also noticed in the example image (Figure S3A-S3B and Table S2).

**Figure 2:**
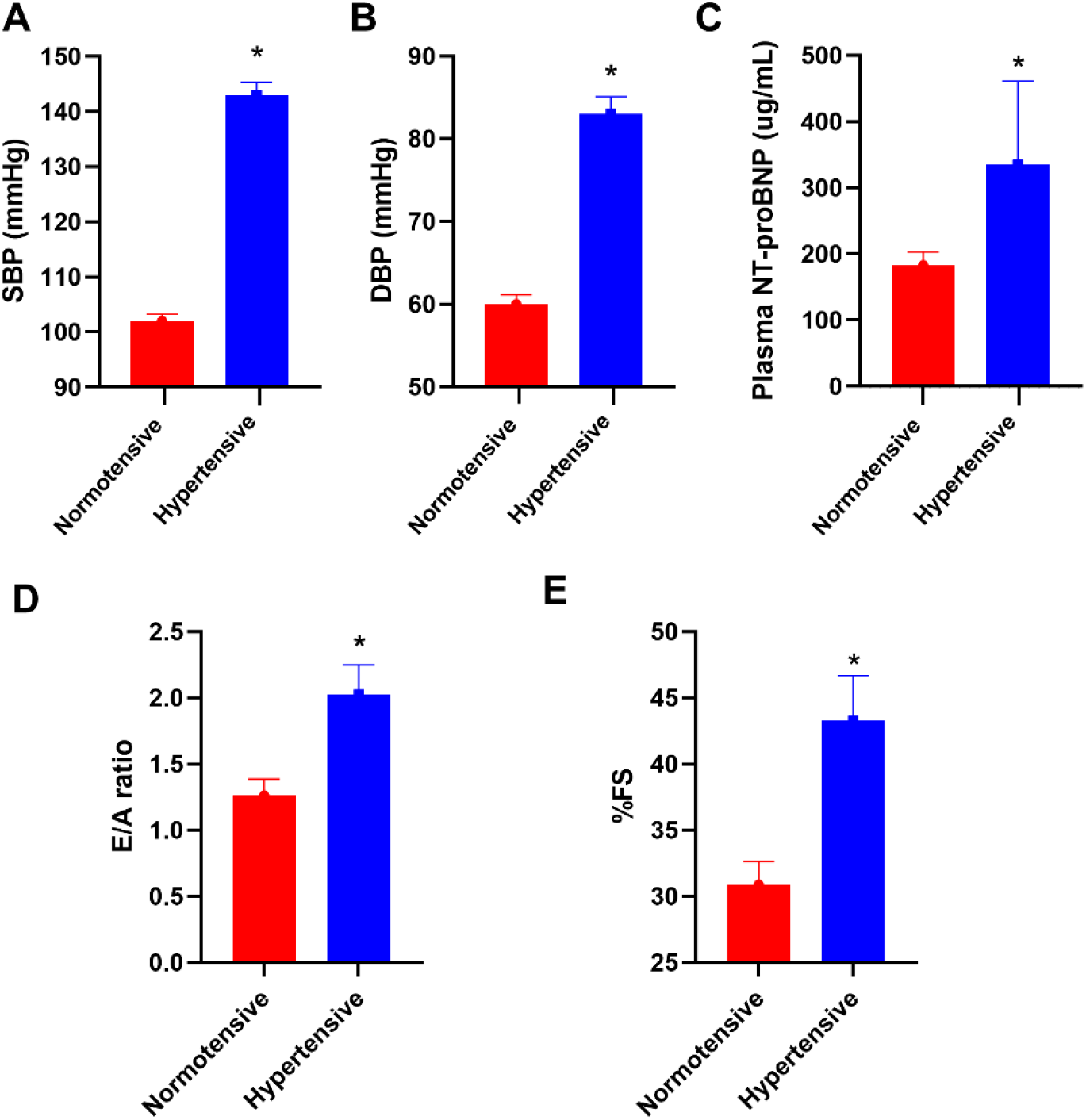
Hypertensive animals had significantly higher levels of (**A**) Higher systolic blood pressure (SBP), (**B**) diastolic blood pressure (DBP), (**C**) plasma concentrations of N-terminal pro-B-type natriuretic peptide (NT-proBNP), (**D**) ratio of the early (E) to late (A) ventricular filling velocities, (**E**) percentage fractional shortening (% FS) compared to normotensive animals in cross-sectional study (study 1). Values are shown as mean ± SEM. **\***p<0.05.

### A strong correlation between MT and blood pressure measures

To understand the effects of metabolic health markers on MT, we performed a Pearson’s correlation analysis. Our correlation analysis showed significant positive correlations existed only between blood pressure measures and MT markers. We observed a significant positive correlation between LBP-1 and sCD14 (r=0.28, n=147, p=0.001), and LBP-1 and DBP (r= 0.19, p=0.02) (Table 1). Both, LBP-1 (r=0.17, p=0.04) and sCD14 (r=0.17, p=0.04) significantly increased with age.

**Table 1:** Correlation analysis between plasma concentrations of lipopolysaccharide-binding protein (LBP-1) in a cross-sectional study (study 1). Abbreviations: R= correlation coefficient, N= sample size, BW= body weight, Waist circum= Waist circumference, FBG= fasting blood glucose, TG= total plasma triglycerides, HDLC= high-density lipoprotein cholesterol, SBP= systolic blood pressure, DBP= diastolic blood pressure. **\***p<0.05.

| Variables | Age | BW | Waist<br>circum | FBG | TG | HDLC | SBP | DBP |
| --- | --- | --- | --- | --- | --- | --- | --- | --- |
| LBP-1 |  |  |  |  |  |  |  |  |
| R | 0.17 | 0.03 | 0.02 | 0.12 | -0.07 | 0.04 | 0.15 | 0.19 |
| N | 147 | 147 | 147 | 147 | 147 | 147 | 147 | 147 |
| P-value | 0.04* | 0.69 | 0.79 | 0.16 | 0.39 | 0.51 | 0.08 | 0.02* |

### Independent effects of hypertension on MT

To determine the relationship between blood pressure and MT measures by their associations with age, we performed a partial regression analysis and multiple predictive regression modeling. The age-adjusted LBP-1 levels remained significantly correlated with SBP (R^2^=0.16, p=0.05) and DBP (R^2^=0.20, p=0.02) (Table 2), while age was not correlated with any of the blood pressure measures. We performed a multivariate analysis to determine the effect of MT on blood pressure regulation. The significant variables in the final model were age (β=0.033, p=0.04) and DBP (β=0.012, p=0.02). This demonstrates the significant and independent effects of blood pressure regulation on MT.

**Table 2:** Age-adjusted correlation between plasma concentrations of lipopolysaccharide-binding protein (LBP-1), an indicator of microbial translocation and systolic blood pressure (SBP) and diastolic blood pressure (DBP) in a cross-sectional study (study 1). R=correlation coefficient.

| **LBP-1** | **Systolic BP** | **Diastolic BP** |
| --- | --- | --- |
| R | 0.16 | 0.20 |
| N | 147 | 147 |
| p-value | 0.05* | 0.02* |

### Study 2: Longitudinal study on hypertension, MT and microbiome

Sixteen adult male rhesus macaques (mean age 5.8±0.5 years) were included in this study. These animals were divided into hypertensive (N=8) and normotensive (N=8), and their metabolic health characteristics at baseline and 27-month time points are given in Table S3. The SBP and DBP in normotensive monkeys consistently showed low levels, while in hypertensive monkeys, these pressure values were higher (repeated measures ANOVA group effect p=0.008) and continued to be different throughout the > 2 year study (SBP p=0.03 and DBP p=0.12 [Figure 3A and 3B]). We observed significantly higher plasma concentrations of NT-proBNP in hypertensive animals at baseline and these also stayed higher at study end (p=0.04). Overall, the corrected means for the hypertensive group from repeated measures of ANOVA showed a significant group effect (p=0.04) for plasma concentrations of NT-proBNP being elevated with hypertension (Figure 3C).

**Figure 3:**
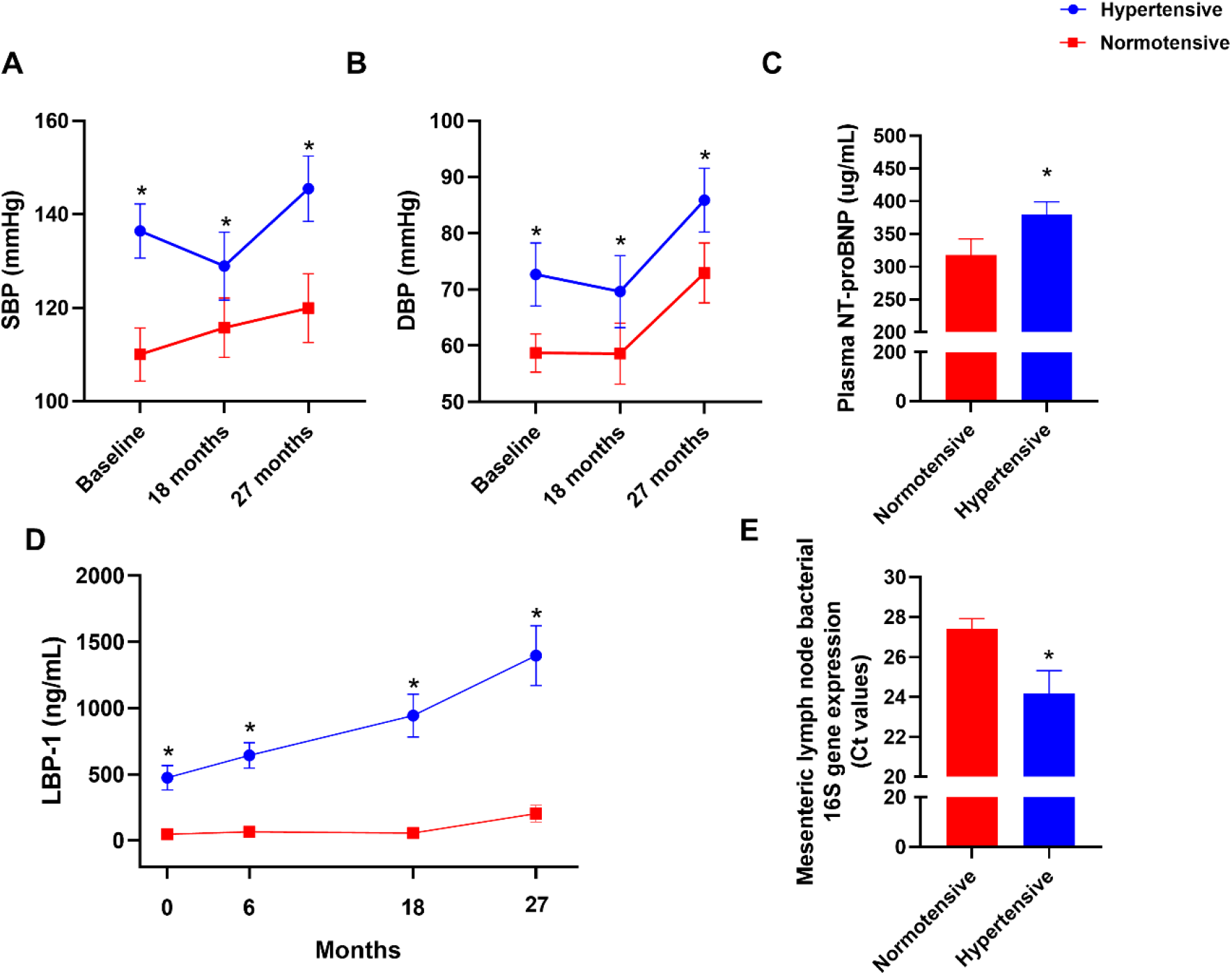
Hypertensive animals had significantly higher levels of (**A**) systolic blood pressure (SBP), (**B**) diastolic blood pressure (DBP), (**C**) plasma concentrations of N-terminal pro-B-type natriuretic peptide (NT-proBNP), (**D**) lipopolysaccharide-binding protein (LBP)-1 and (**E**) lower 16S bacterial gene expression in mesenteric lymph nodes compared to normotensive animals in longitudinal study (study 2), where Ct value for 16S; lower = higher gene expression. Values are shown as mean ± SEM. **\***p<0.05.

### Higher MT in the hypertensive group

At baseline, the individual MT biomarker (LBP-1 concentrations) was higher in hypertensive animals, with >14-fold difference between these two groups (46 vs. 668 ng/mL, respectively) and these levels progressively and dramatically increased over two years, compared to the normotensive animals (Figure 3D). Further, we observed a higher level of 16S bacterial gene presence in extra-intestinal tissues, which confirms higher MT and barrier dysfunction in hypertensive animals compared to normotensive counterparts (Figure 3E).

### Fecal microbiome dysbiosis in the hypertensive group

For the fecal microbiome samples, there was no difference in the number of microbial sequences per samples, or the alpha (Figure S3A) and beta diversity profiles (Figure S3B and S3C); however, taxonomic changes were observed based on blood pressure status. At the phylum level, Firmicutes, Bacteroidetes, Proteobacteria and Actinobacteria were most predominantly present in both the groups. At baseline, the percentage counts of the opportunist bacterial order Clostridiales were higher in the hypertensive group (Figure S3D). During the follow-up period at 12 months and 27 months, the percentage counts of the beneficial bacterial family, *Lactobacillaceae* (Figure 4A) were decreased, and the percentage counts of the opportunistic family such as *Ruminococcaceae* (Figure 4B) were increased in the hypertensive group compared to normotensive counterparts.

**Figure 4:**
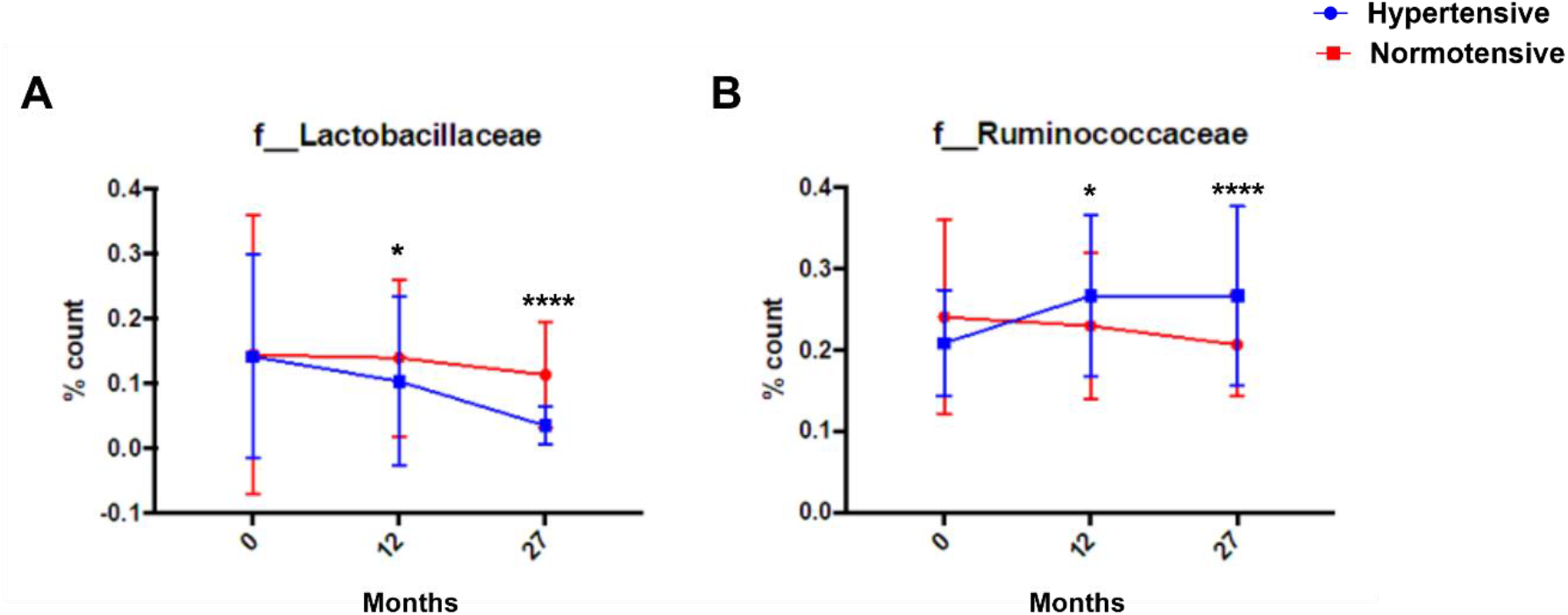
Development of dysbiosis in fecal microbiomes in hypertensive animals in longitudinal study (study 2). **A**. The percentage counts of families *Lactobacillaceae* **B**. *Ruminococcaceae* (order Clostridiales) from baseline to 27 months. Values are shown as mean ± SEM. **\***p<0.05, **** < 0.0001.

## Discussion

To the best of our knowledge, this is the first study to examine the associations between gut barrier dysfunction and dysbiosis with hypertension in a nonhuman or human primate model that includes a longitudinal component to the study design, and thus has high translational relevance to human health. In the present study, we showed that hypertensive animals had higher concentrations of plasma LBP-1 (a MT biomarker) and dysbiotic shifts in the gut microbiome compared to normotensive animals, suggesting that hypertension leads to MT and dysbiosis. Specifically, our major findings are that (i) hypertensive animals had higher MT, measured by plasma LBP-1 concentration and extra-intestinal bacteria abundance when compared to normotensive animals; (ii) increases in plasma concentration of LBP-1 are positively correlated with DBP, independent of age and other health markers, suggesting specificity of the independent effect of hypertension and; (iii) in longitudinal analyses of environmentally controlled individuals, hypertensive animals had dysbiotic remodeling of their microbiome towards a less healthy profile. These findings suggest that hypertension is a contributor to MT and dysbiosis and may contribute to poor health in ways not considered before, and points potentially to another important reason for an early diagnosis and treatment of hypertension. As MT and dysbiosis drive inflammatory processes along with hypertension-related changes in tissue perfusion and function, we describe a novel connection between the gut and heart.

The findings of our cross-sectional study (study 1) are consistent with a few animal studies and recent clinical studies, which showed the associations between hypertension and gut barrier dysfunction.^7, 10, 12, 22^ Nonetheless, the aforementioned studies were cross-sectional and were designed to examine associations and not to determine cause-and-effect relationships. Besides, the gut microbiota is considered as a regulator of the gut barrier function, and studies have shown dysbiotic changes in the gut microbial composition with hypertension.^6, 23^ However, no study has investigated whether dysbiosis precede, accompany, or result from hypertension. To address these gaps, we performed a longitudinal study (study 2) to identify the gut microbiota changes in hypertension.

We have previously demonstrated LBP-1 as a reliable marker for MT and its positive association with age-related metabolic health decline in NHPs and humans.^3–5, 15^ More recently, a Japanese-based population longitudinal study showed that higher LBP-1 levels are associated with an increased risk of the CVD development.^10^ However, no studies have investigated the role of MT in hypertension, which is a major contributor to CVD. Our cross-sectional study (study 1) found the evidence to support our hypothesis, as mentioned in the introduction section, that MT is associated with hypertension, an indicator of gut barrier dysfunction. A dysfunctional gut barrier promotes lipopolysaccharide (measure by LBP-1 concentrations) of the gut microbiota to the host, thereby inducing host inflammation and, finally, impacting cardiovascular health.^9, 10, 12^ The combination of hypertension and MT was also observed alongside higher NT-proBNP and impaired diastolic function measure by echocardiogram, which relates to the risk of heart failure.^24, 25^ NT-proBNP has recently attracted a lot of attention as a biomarker for heart function. The parent BNP peptide is synthesized by atrial and ventricular cardiomyocytes, and exert potent diuretic, natriuretic, vasorelaxant and favorable metabolic effects, which are mediated via guanyl cyclase-A. BNP suppresses the renin-angiotensin pathway and sympathetic activity. ^26, 27^ Collectively, the parent protein should exert positive effects, however, it has been found in clinical trials to be useful both diagnostically and prognostically within the range of concentrations that we report here measured from at-risk monkeys. ^28^

Our observed association between hypertension and MT was independent of age. Although aging is associated with worsening gut barrier function, our observed association between hypertension and age occurred even when age was accounted for. Previously, we showed that age-related increases in MT and western diet, either alone or in combination, led to a decline in metabolic health.^3, 5, 15^ However, in the present study, LBP-1 levels were not associated with any components of metabolic syndrome, which could be related to the healthy diet consumed by these animals.

These findings demonstrate that the association between hypertension and MT is partially driven by aging, but hypertension is an independent effect that worsens the overall health.

To confirm the involvement of the gut microbiome, we collected fecal samples over a two-year period (baseline, 12 and 27-month time points) and investigated microbiome changes using 16S rRNA gene sequencing technology to identify gut microbiota changes between hypertensive versus normotensive animals. Consistent with our hypothesis, the results from our longitudinal study (study 2) demonstrated that the hypertension drives echocardiographic markers that suggest higher risk of heart failure and drove gut barrier dysfunction, which led to the fecal microbiome perturbations in hypertensive animal. Moreover, for the first time, we found that dysbiosis did manifest at baseline, but was more evidently discerned at 12-month and remained dysbiotic at the 27-month time point in hypertensive animals compared to normotensive animals. Particularly, at baseline, noticeable changes were increases in the opportunistic bacterial order, Clostridiales in hypertensive animals compared to normotensive animals. However, at the 12-month and 27-month time points, at the family level, we found increases in *Ruminococcaceae* (Clostridiales order) and decreases in *Lactobacillaceae* families in the hypertensive group compared to the normotensive group. Such alterations could represent robust dysbiotic trends in the microbial community, suggesting that the dysbiotic changes are a consequence of hypertension.^6, 23^

Several mechanisms might explain why hypertension can induce gut barrier dysfunction and dysbiosis. Firstly, animal studies have shown that hypertension triggered changes in arterioles (a part of microcirculatory system), which decreased intestinal blood flow.^12, 13^ In addition, from recent human studies, we know that hypertension induces vascular pathologies such as arterial stiffness, which reduce intestinal perfusion.^23, 29^ Decreased blood flow and intestinal perfusion dampen the nutrient supply and absorption in the gut, causing gut mucosal damage and barrier dysfunction leading to dysbiosis.^23, 30^ Secondly, gut microbial-derived metabolites such as short-chained fatty acids (SCFAs) are known to energize the gut epithelium, enhance gut permeability, and regulate blood pressure.^9, 23, 31^ Particularly, SCFAs may regulate blood pressure by acting on the G protein-coupled and olfactory receptors, expressed in the kidney and smooth muscle cells of blood vessels respectively.^32^ Although a few members of the *Ruminococcaceae* family are recognized as SCFA producers in healthy conditions, they are inversely correlated with gut permeability and overabundant in some gut inflammatory disorders.^33^ In contrast, *Lactobacillaceae* family members, particularly the taxa, *Lactobacillus* and *Bifidobacterium*, are known for their beneficial effects via an increase in SCFA production.^31, 33^ Thus, microbiome-induced SCFA-led alterations could be one of the main mechanisms of the hypertension condition, as observed in our study. Although there is evidence regarding the significant role of gut microbiome produced SCFA in modulating blood pressure, the role of SCFA in hypertension warrants further research.

The strengths of our study include the prospective cohort study design, larger sample size and ideal experimental conditions for follow-up over two years, coupled with an accurate diagnosis of hypertension and MT based on available clinical information. Moreover, this could be the first longitudinal study to show the gut microbiome shifts are resulting from hypertension. We also address repeatability and generalizability of the relationship between hypertension and gut barrier dysfunction via the use of two separate nonhuman primate cohorts and species, which show consistent results. However, there are a few limitations of the present study. A subset of animals from the study 2 received TBI, however we found no significant differences in their LBP, BP measures and microbiome (Figure S4). Plasma biomarkers only indirectly reflect gut barrier dysfunction. However, these biomarkers act as potential predictors for screening and diagnosis of the disease condition and we confirmed barrier dysfunction by detection of higher bacterial genome levels in lymph nodes draining the colon.^34^ While providing novel information about microbial signatures of hypertension, possibly more robust work is needed at the species level to understand the alterations in microbial metabolism and inflammation and their contributions to hypertension. Understanding the host-microbiome-metabolic axis will enable us to gain insights into diagnosis and develop a microbiome-related therapeutic intervention to manage blood pressure in hypertensive patients. To this end, probiotics are natural and safe modulators of the gut microbiome.^9, 33^ Although bacterial probiotics, are attractive options, it is still unclear if they would benefit hypertensive patients.^35^ Instead, engineered bacteria probiotics, such as *Lactobacillus* spp. to target specific tissues and cells rather than the whole body might be new modalities to develop a therapeutic approach on a clinical scale.

### Perspectives

The present study indicates that hypertension is associated with MT and gut barrier dysfunction and the risk of heart failure in spontaneous hypertensive primates (study 1). Further, hypertension and gut barrier dysfunction, either alone or in combination, triggered the gut microbial dysbiosis consistently over two years (study 2). Our data indicated support for the role of the novel “heart-gut” axis by showing a high risk of heart failure, elevated NT-proBNP concentrations as well as cardiac pathology (echocardiography) during hypertension, which led to gut barrier dysfunction and dysbiosis.^36^ Thus, the interaction between gut microbiota and host can be applied as a potential therapeutic option to help overcome the global burden of hypertension and CVD. Additional studies are required to determine the detailed microbial metabolism and inflammatory mechanisms of the gut barrier dysfunction and dysbiosis linked to hypertension.

## Acknowledgments

We would like to thank the animal care, veterinary, and research staff at the Wake Forest School of Medicine Clarkson Campus, UNC Microbiome Core Facility (the Center for Gastrointestinal Biology and Disease (CGIBD P30 DK034987) and the UNC Nutrition Obesity Research Center (NORC P30 DK056350) as well as the NHPs themselves.

## References

1. Genua F, Raghunathan V, Jenab M, Gallagher WM, Hughes DJ. The role of gut barrier dysfunction and microbiome dysbiosis in colorectal cancer development. Front Oncol. 2021;11:626349–626349

2. Peery AF, Crockett SD, Murphy CC, Lund JL, Dellon ES, Williams JL, Jensen ET, Shaheen NJ, Barritt AS, Lieber SR, Kochar B, Barnes EL, Fan YC, Pate V, Galanko J, Baron TH, Sandler RS. Burden and cost of gastrointestinal, liver, and pancreatic diseases in the united states: Update 2018. Gastroenterology. 2019;156:254–272.e211

3. Kavanagh K, Hsu F-C, Davis AT, Kritchevsky SB, Rejeski WJ, Kim S. Biomarkers of leaky gut are related to inflammation and reduced physical function in older adults with cardiometabolic disease and mobility limitations. Geroscience. 2019;41:923–933

4. Vemuri R, Sherrill C, Davis MA, Kavanagh K. Age-related colonic mucosal microbiome community shifts in monkeys. The Journals of Gerontology: Series A. 2020

5. Wilson QN, Wells M, Davis AT, Sherrill C, Tsilimigras MCB, Jones RB, Fodor AA, Kavanagh K. Greater microbial translocation and vulnerability to metabolic disease in healthy aged female monkeys. Scientific reports. 2018;8:1–10

6. Kim S, Goel R, Kumar A, Qi Y, Lobaton G, Hosaka K, Mohammed M, Handberg EM, Richards EM, Pepine CJ. Imbalance of gut microbiome and intestinal epithelial barrier dysfunction in patients with high blood pressure. Clinical science. 2018;132:701–718

7. Drolia R, Bhunia AK. Crossing the intestinal barrier via listeria adhesion protein and internalin a. Trends in microbiology. 2019;27:408–425

8. Yu LC-H. Microbiota dysbiosis and barrier dysfunction in inflammatory bowel disease and colorectal cancers: Exploring a common ground hypothesis. Journal of biomedical science. 2018;25:1–14

9. Vemuri R, Gundamaraju R, Shastri MD, Shukla SD, Kalpurath K, Ball M, Tristram S, Shankar EM, Ahuja K, Eri R. Gut microbial changes, interactions, and their implications on human lifecycle: An ageing perspective. BioMed research international. 2018;2018

10. Asada M, Oishi E, Sakata S, Hata J, Yoshida D, Honda T, Furuta Y, Shibata M, Suzuki K, Watanabe H. Serum lipopolysaccharide-binding protein levels and the incidence of cardiovascular disease in a general japanese population: The hisayama study. Journal of the American Heart Association, 2019;8:e013628

11. Lepper PM, Schumann C, Triantafilou K, Rasche FM, Schuster T, Frank H, Schneider EM, Triantafilou M, von Eynatten M. Association of lipopolysaccharide-binding protein and coronary artery disease in men. Journal of the American College of Cardiology. 2007;50:25–31

12. Jaworska K, Huc T, Samborowska E, Dobrowolski L, Bielinska K, Gawlak M, Ufnal M. Hypertension in rats is associated with an increased permeability of the colon to tma, a gut bacteria metabolite. PloS one, 2017;12:e0189310

13. Santisteban MM, Qi Y, Zubcevic J, Kim S, Yang T, Shenoy V, Cole-Jeffrey CT, Lobaton GO, Stewart DC, Rubiano A. Hypertension-linked pathophysiological alterations in the gut. Circulation research. 2017;120:312–323

14. Lerman LO, Kurtz TW, Touyz RM, Ellison DH, Chade AR, Crowley SD, Mattson DL, Mullins JJ, Osborn J, Eirin A. Animal models of hypertension: A scientific statement from the american heart association. Hypertension, 2019;73:e87–e120

15. Mitchell EL, Davis AT, Brass K, Dendinger M, Barner R, Gharaibeh R, Fodor AA, Kavanagh K. Reduced intestinal motility, mucosal barrier function, and inflammation in aged monkeys. The journal of nutrition, health & aging. 2017;21:354–361

16. Kavanagh K, Jones KL, Sawyer J, Kelley K, Carr JJ, Wagner JD, Rudel LL. Trans fat diet induces abdominal obesity and changes in insulin sensitivity in monkeys. Obesity (Silver Spring). 2007;15:1675–1684

17. Kaneko H, Itoh H, Yotsumoto H, Kiriyama H, Kamon T, Fujiu K, Morita K, Michihata N, Jo T, Takeda N. Association of isolated diastolic hypertension based on the cutoff value in the 2017 american college of cardiology/american heart association blood pressure guidelines with subsequent cardiovascular events in the general population. Journal of the American Heart Association, 2020;9:e017963

18. Brownlee RD, Kass PH, Sammak RL. Blood pressure reference intervals for ketamine-sedated rhesus macaques (macaca mulatta). J Am Assoc Lab Anim Sci. 2020;59:24–29

19. Bacarella N, Ruggiero A, Davis AT, Uberseder B, Davis MA, Bracy DP, Wasserman DH, Cline JM, Sherrill C, Kavanagh K. Whole body irradiation induces diabetes and adipose insulin resistance in nonhuman primates. International Journal of Radiation Oncology*Biology*Physics. 2020;106:878–886

20. DeBo RJ, Lees CJ, Dugan GO, Caudell DL, Michalson KT, Hanbury DB, Kavanagh K, Cline JM, Register TC. Late effects of total-body gamma irradiation on cardiac structure and function in male rhesus macaques. Radiat Res. 2016;186:55–64

21. Guo H, Chou W-C, Lai Y, Liang K, Tam JW, Brickey WJ, Chen L, Montgomery ND, Li X, Bohannon LM. Multi-omics analyses of radiation survivors identify radioprotective microbes and metabolites. Science. 2020;370

22. Li C, Xiao P, Da Lin H-JZ, Zhang R, Zhao Z-g, He X-X. Risk factors for intestinal barrier impairment in patients with essential hypertension. Frontiers in medicine. 2020;7

23. Silveira-Nunes G, Durso DF, Cunha EHM, Maioli TU, Vieira AT, Speziali E, Corrêa-Oliveira R, Martins-Filho OA, Teixeira-Carvalho A, Franceschi C. Hypertension is associated with intestinal microbiota dysbiosis and inflammation in a brazilian population. Frontiers in pharmacology, 2020;11:258

24. Rørth R, Jhund PS, Yilmaz MB, Kristensen SL, Welsh P, Desai AS, Køber L, Prescott MF, Rouleau JL, Solomon SD. Comparison of bnp and nt-probnp in patients with heart failure and reduced ejection fraction. Circulation: Heart Failure, 2020;13:e006541

25. Salah K, Stienen S, Pinto YM, Eurlings LW, Metra M, Bayes-Genis A, Verdiani V, Tijssen JGP, Kok WE. Prognosis and nt-probnp in heart failure patients with preserved versus reduced ejection fraction. Heart. 2019;105:1182–1189

26. Buglioni A, Cannone V, Cataliotti A, Sangaralingham SJ, Heublein DM, Scott CG, Bailey KR, Rodeheffer RJ, Dessi-Fulgheri P, Sarzani R, Burnett JC, Jr. Circulating aldosterone and natriuretic peptides in the general community: Relationship to cardiorenal and metabolic disease. Hypertension. 2015;65:45–53

27. Calhoun DA, Jones D, Textor S, Goff DC, Murphy TP, Toto RD, White A, Cushman WC, White W, Sica D, Ferdinand K, Giles TD, Falkner B, Carey RM, American Heart Association Professional Education C. Resistant hypertension: Diagnosis, evaluation, and treatment: A scientific statement from the american heart association professional education committee of the council for high blood pressure research. Circulation, 2008;117:e510–526

28. Maisel AS, Duran JM, Wettersten N. Natriuretic peptides in heart failure: Atrial and b-type natriuretic peptides. Heart Fail Clin. 2018;14:13–25

29. Nakanishi R, Baskaran L, Gransar H, Budoff MJ, Achenbach S, Al-Mallah M, Cademartiri F, Callister TQ, Chang H-J, Chinnaiyan K. Relationship of hypertension to coronary atherosclerosis and cardiac events in patients with coronary computed tomographic angiography. Hypertension. 2017;70:293–299

30. Granger DN, Holm L, Kvietys P. The gastrointestinal circulation: Physiology and pathophysiology. Comprehensive Physiology. 2015;5:1541–1583

31. Vemuri R, Gundamaraju R, Shinde T, Perera AP, Basheer W, Southam B, Gondalia SV, Karpe AV, Beale DJ, Tristram S. Lactobacillus acidophilus dds-1 modulates intestinal-specific microbiota, short-chain fatty acid and immunological profiles in aging mice. Nutrients, 2019;11:1297

32. Pluznick J. A novel scfa receptor, the microbiota, and blood pressure regulation. Gut microbes. 2014;5:202–207

33. Vemuri RC, Gundamaraju R, Shinde T, Eri R. Therapeutic interventions for gut dysbiosis and related disorders in the elderly: Antibiotics, probiotics or faecal microbiota transplantation? Beneficial microbes. 2017;8:179–192

34. Ellison S, Abdulrahim JW, Kwee LC, Bihlmeyer NA, Pagidipati N, McGarrah R, Bain JR, Kraus WE, Shah SH. Novel plasma biomarkers improve discrimination of metabolic health independent of weight. Scientific reports. 2020;10:1–9

35. Jama HA, Kaye DM, Marques FZ. The gut microbiota and blood pressure in experimental models. Current opinion in nephrology and hypertension. 2019;28:97–104

36. Forkosh E, Ilan Y. The heart-gut axis: New target for atherosclerosis and congestive heart failure therapy. Open heart, 2019;6:e000993

