## Supplementary material for "Hypertension drives microbial translocation and shifts in the fecal microbiome of non-human primates": Table S1: Vemuri et al. Hypertension Data suppl.docx

DATA SUPPLEMENT

| Variables | N | Mean | Median | Minimum | Maximum | SD | SEM |
| --- | --- | --- | --- | --- | --- | --- | --- |
| Age (yrs.)  Male  Female | 153  24  129 | 16 | 17 | 6.4 | 28 | 5 | 0.4 |
| BW (kg) | 153 | 6 | 6 | 3.5 | 11 | 1 | 0.1 |
| Waist circum (cm) | 153 | 36 | 36 | 25 | 51 | 5 | 0.4 |
| FBG (mg/dL) | 153 | 82 | 71 | 47 | 429 | 43 | 3.5 |
| Hemoglobin (Hb)A1c | 153 | 4 | 4 | 3.9 | 10 | 1 | 0.1 |
| SBP (mmHg) | 153 | 120 | 115 | 74.3 | 208 | 25 | 2 |
| DBP (mmHg) | 153 | 70 | 67 | 33 | 128 | 18 | 1.5 |
| HDL-C (mg/dL) | 153 | 113 | 120 | 9.7 | 264 | 54 | 4.3 |
| TPC (mg/dL) | 153 | 64 | 58 | 25.1 | 279 | 31 | 2.5 |
| LBP-1 (ng/mL) | 147 | 155 | 59 | 9.8 | 1276 | 222 | 18.3 |
| sCD14 (ng/L) | 148 | 4109 | 3747 | 617.6 | 10137 | 2014 | 165.6 |

**Table S1**: Basic demographic and metabolic details in the cross-sectional study (study 1). Abbreviations: BW= body weight, Waist circum= Waist circumference, FBG= fasting blood glucose, SBP= systolic blood pressure, DBP= diastolic blood pressure, HDLC= high density lipoprotein cholesterol, TPC=total plasma cholesterol, LBP-1= lipopolysaccharide binding protein-1, sCD14= Soluble CD14, SD=standard deviation, SEM=standard error of the mean.

|  | **Normotensive** | | | | **Hypertensive** | | | |
| --- | --- | --- | --- | --- | --- | --- | --- | --- |
| **Variables** | Means | N | SD | SEM | Means | N | SD | SEM |
| A/E (ratios) | 0.862 | 10 | 0.275 | 0.086 | 0.552 | 11 | 0.188 | 0.056 |
| Aa(l) m/s | 0.060 | 10 | 0.010 | 0.003 | 0.051 | 11 | 0.016 | 0.004 |
| AV PGmax (mmHg) | 2.828 | 10 | 0.989 | 0.313 | 3.555 | 11 | 2.242 | 0.676 |
| Aa(s) m/s | 0.055 | 10 | 0.015 | 0.004 | 0.049 | 10 | 0.017 | 0.005 |
| AV PGmean (mmHg) | 1.200 | 10 | 0.379 | 0.119 | 1.704 | 11 | 1.040 | 0.313 |
| AV Vmax (m/s) | 0.826 | 10 | 0.165 | 0.052 | 0.906 | 11 | 0.272 | 0.082 |
| AV Vmean (m/s) | 0.488 | 10 | 0.075 | 0.023 | 0.580 | 11 | 0.176 | 0.053 |
| AV VTI (cm) | 8.433 | 10 | 2.563 | 0.810 | 10.404 | 11 | 2.753 | 0.830 |
| Dec Slope (m/s2) | 7.720 | 10 | 2.505 | 0.792 | 9.373 | 11 | 3.786 | 1.141 |
| Dec Time (ms) | 92.547 | 10 | 24.490 | 7.744 | 83.934 | 11 | 23.532 | 7.095 |
| E/Ea(l) (ratio) | 8.017 | 10 | 1.970 | 0.623 | 8.529 | 11 | 2.241 | 0.675 |
| Ea(l) (m/s) | 0.086 | 10 | 0.022 | 0.007 | 0.088 | 11 | 0.023 | 0.007 |
| Ea(s) (m/s) | 0.077 | 10 | 0.012 | 0.004 | 0.073 | 10 | 0.017 | 0.005 |
| Ea/Aa(l) (ratio) | 1.468 | 10 | 0.423 | 0.133 | 1.907 | 11 | 0.894 | 0.269 |
| Ea/Aa(s) ratio | 1.501 | 10 | 0.498 | 0.157 | 1.578 | 10 | 0.550 | 0.173 |
| EDV (A4C) mL | 4.899 | 10 | 1.001 | 0.316 | 4.023 | 11 | 2.275 | 0.686 |
| EF (A4C) % | 49.574 | 10 | 9.045 | 2.860 | 44.690 | 11 | 14.332 | 4.321 |
| ESV (A4C) mL | 2.433 | 10 | 0.511 | 0.161 | 2.295 | 11 | 1.482 | 0.447 |
| LA Vol (A2C) mL | 0.930 | 10 | 0.443 | 0.140 | 0.890 | 11 | 0.441 | 0.133 |
| LA Vol (A4C) mL | 1.579 | 10 | 0.606 | 0.191 | 1.505 | 10 | 0.995 | 0.314 |
| LA Vol (BP) mL | 1.345 | 10 | 0.594 | 0.187 | 1.192 | 10 | 0.609 | 0.192 |
| LAA (A2C) cm2 | 1.146 | 10 | 0.356 | 0.112 | 1.092 | 10 | 0.347 | 0.109 |
| LAA (A4C) cm2 | 1.688 | 10 | 0.391 | 0.123 | 1.590 | 11 | 0.602 | 0.181 |
| LVOT diam (mm) | 7.725 | 10 | 0.787 | 0.249 | 7.510 | 12 | 1.110 | 0.320 |
| MV A pt (m/s) | 0.538 | 10 | 0.060 | 0.019 | 0.396 | 11 | 0.143 | 0.043 |
| MV E pt (m/s) | 0.668 | 10 | 0.162 | 0.051 | 0.733 | 11 | 0.180 | 0.054 |
| SV (A4C) mL | 2.465 | 10 | 0.766 | 0.242 | 1.728 | 11 | 1.051 | 0.317 |
| IVSd (cm) | 0.490 | 10 | 0.120 | 0.038 | 0.512 | 12 | 0.130 | 0.037 |
| LVIDd (mm/m2) | 1.761 | 10 | 0.370 | 0.117 | 1.665 | 12 | 0.380 | 0.109 |
| LVPWd (mm/m2) | 0.492 | 10 | 0.180 | 0.057 | 0.493 | 12 | 0.195 | 0.056 |
| IVSs (cm) | 0.519 | 10 | 0.182 | 0.057 | 0.611 | 12 | 0.187 | 0.054 |
| LVPWs (cm) | 0.673 | 10 | 0.122 | 0.038 | 0.833 | 12 | 0.235 | 0.068 |
| EDV teich (mL) | 9.900 | 10 | 4.867 | 1.539 | 8.689 | 12 | 4.424 | 1.277 |
| ESV teich (mL) | 3.663 | 10 | 1.651 | 0.522 | 2.250 | 12 | 2.077 | 0.599 |
| EF teich (mL) | 61.608 | 10 | 7.934 | 2.509 | 75.853 | 12 | 14.408 | 4.159 |
| SV teich (mL) | 6.237 | 10 | 3.370 | 1.065 | 6.439 | 12 | 2.893 | 0.835 |

**Table S2:** Difference between echocardiogram parameters in the hypertensive group compared to the normotensive group in the cross-sectional study (study 1). A/E= Atrial contraction/early diastole, Aa(l) & Aa(s)= atrial enlargement, AV PGmax= aortic valve maximal pressure gradient, AV PGmean= aortic valve mean pressure gradient, AV Vmax= aortic valve maximum velocity, AV Vmean= aortic valve mean velocity, AV VTI= aortic valve area, Dec Slope= deceleration slope, Dec Time= deceleration time, E/Ea= early diastolic mitral annular tissue velocity, EDV= end-diastolic volume, EF= ejection fraction, ESV= end-systolic volume, LA = Left atrial, LVOT= left ventricular outflow tract, MV= mitral valve area, IVSd and IVSs= Interventricular septal end diastole and end systole, LVPWd= Left ventricular posterior wall thickness at end-diastole, LVPW=left ventricle posterior wall, teich= Teichholz method, ESV= End-systolic volume, EF= ejection fraction and SV= stroke volume. SD=standard deviation, SEM=standard error of the mean.

|  | **Baseline** | | **Study End (27 months)** | |
| --- | --- | --- | --- | --- |
| **Variables** | **Normotensive (n=8)** | **Hypertensive (n=8)** | **Normotensive (n=8)** | **Hypertensive (n=8)** |
| Age (years) | 5.02 (0.16) | 5.04 (0.16) |  |  |
| BW (kg) | 9.08 (0.62) | 10.4 (0.82) | 15.7 (1.08) | 16.7 (1.28) |
| Body Fat (%) | 15.4 (1.27) | 18.8 (1.94) | 24.6 (3.00) | 30.0 (2.32) |
| Waist circumference (cm) | 36.4 (1.71) | 38.8 (2.95) | 47.3 (3.69) | 51.7 (2.56) |
| FBG (mg/dL) | 64.5 (3.17) | 67.5 (2.96) | 76.1 (23.8) | 87.5 (17.5) |
| A1c (%) | 4.71 (0.08) | 4.79 (0.11) | 5.83 (0.75) | 5.40 (0.44) |
| TPC (mg/dL) | 161 (15.6) | 173 (7.67) | 204 (18.9) | 179 (18.4) |
| HDLC (mg/dL) | 98.4 (9.58) | 115 (12.3) | 120 (9.74) | 113 (7.95) |
| TG (mg/dL) | 37.0 (11.9) | 39.8 (5.55) | 39.9 (9.24) | 57.4 (18.4) |

**Table S3:** Basic demographic and metabolic details of the longitudinal study (study 2) at baseline and end of study (27 months). The values are shown as the mean ± SEM. Abbreviations: BW= body weight, TPC=total plasma cholesterol, TG=total plasma triglyceride, FBG=fasting glucose, HDLC= high density lipoprotein cholesterol BF= body fat by DEXA scanning, A1c =glycosylated hemoglobin A1c.

|  | **Baseline** | | | | **End of study** | | | |
| --- | --- | --- | --- | --- | --- | --- | --- | --- |
|  | **Control** | | **TBI** | | **Control** | | **TBI** | |
| **Variables** | **N** | **Mean(SEM)** | **N** | **Mean(SEM)** | **N** | **Mean(SEM)** | **N** | **Mean(SEM)** |
| BW | 6 | 9.56 (0.58) | 10 | 9.81 (0.78) | 6 | 18.10833 (1.420) | 10 | 15.09 (0.854) |
| FBG | 6 | 6.88 (3.19) | 10 | 65.3 (2.93) | 6 | 69.83 (12.42) | 10 | 89 (21.97) |
| A1c | 6 | 4.91 (0.09) | 10 | 4.65 (0.07) | 6 | 4.91 (0.03) | 10 | 6.03 (0.65) |
| TPC | 6 | 182.8 (17.76) | 10 | 157.3 (7.91) | 6 | 185.5 (24.28) | 10 | 194.5 (16.11) |
| HDLC | 3 | 95.66 (16.47) | 5 | 110.2 (7.90) | 3 | 118.99 (7.65) | 5 | 114.5 (8.77) |
| TG | 6 | 47.5 (14.58) | 10 | 32.9 (5.30) | 6 | 33 (3.66) | 10 | 58 (15.71) |
| BNP | 4 | 0.22 (0.03) | 9 | 0.23 (0.22) | 4 | 0.23 (0.03) | 9 | 0.23 (0.01) |

**Table S3:** Basic demographic and metabolic details at baseline and end of study (27 months) of irradiated cohort from the longitudinal study (study 2). The values are shown as the mean ± SEM. Abbreviations: BW= body weight, TPC=total plasma cholesterol, TG=total plasma triglyceride, FBG=fasting glucose, HDLC= high density lipoprotein cholesterol, A1c =glycosylated hemoglobin A1c, BNP= B-type natriuretic peptide, TBI= total body irradiation. SEM=standard error.

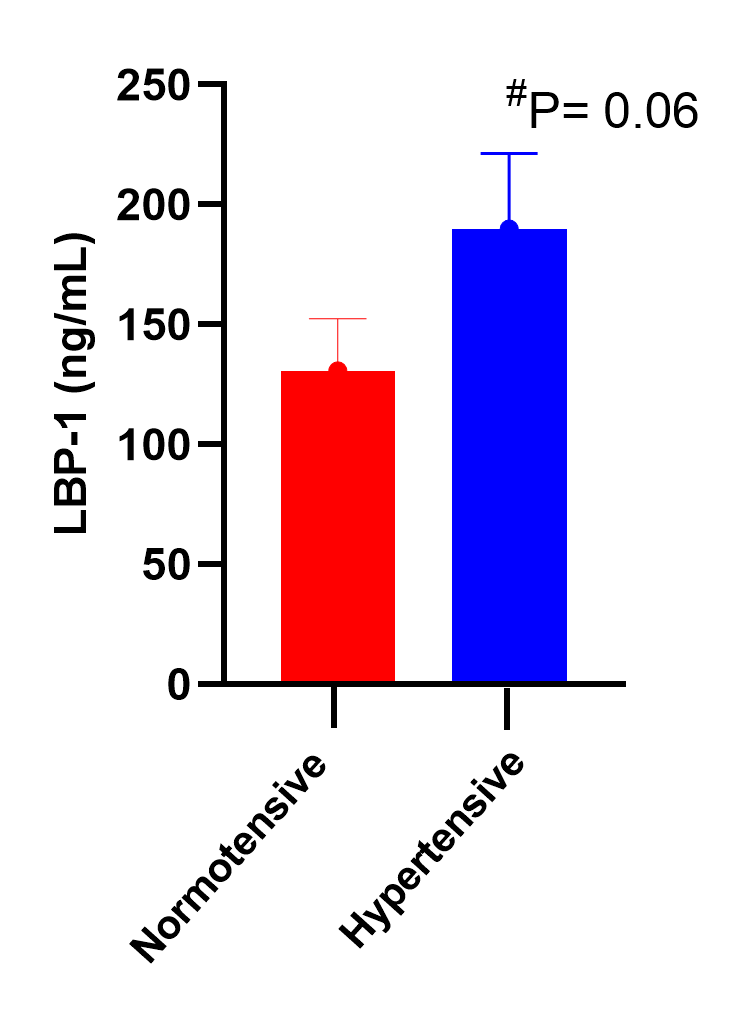

**Figure S1:** In a larger cross sectional study (study 1) of vervet monkeys (N=147), we detected higher lipopolysaccharide-binding protein (LBP-1) in hypertensive (N=61) monkeys compared to the normotensive monkeys (N=86; p=0.06). Values are shown as mean ± standard error.

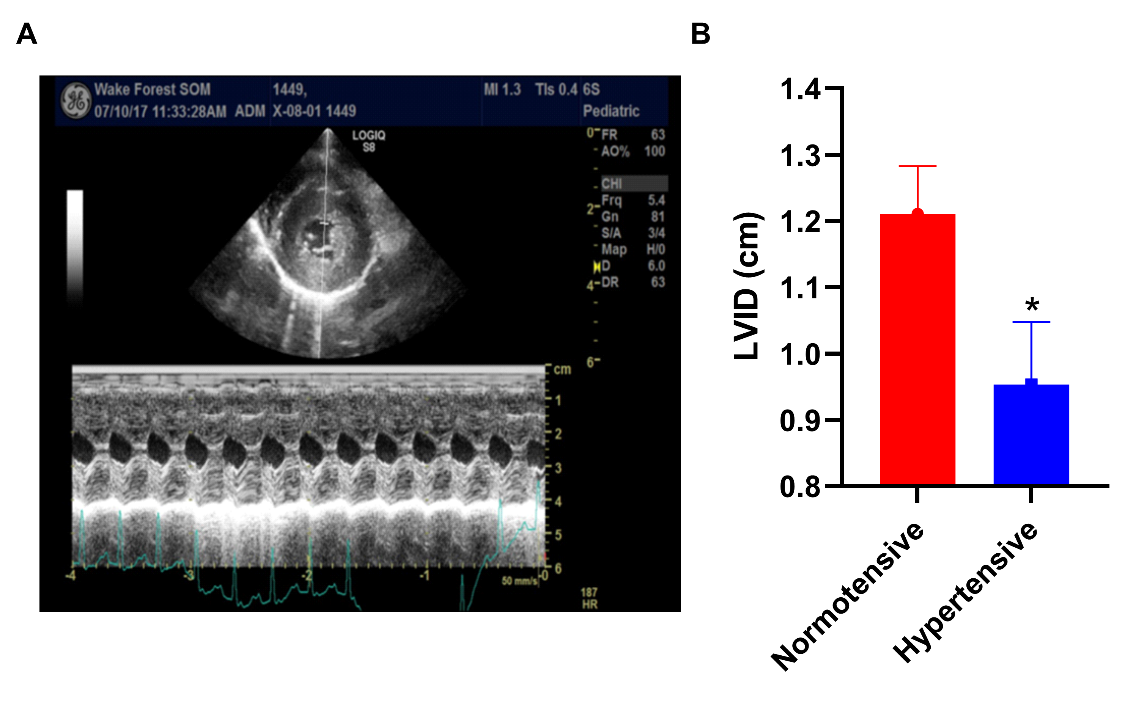

**Figure S2:** An example image from a hypertensive monkey showing dramatic remodeling (**A**) and (**B**) almost no left ventricular internal diameter at systole (LVIDs) in the longitudinal study (study 2).

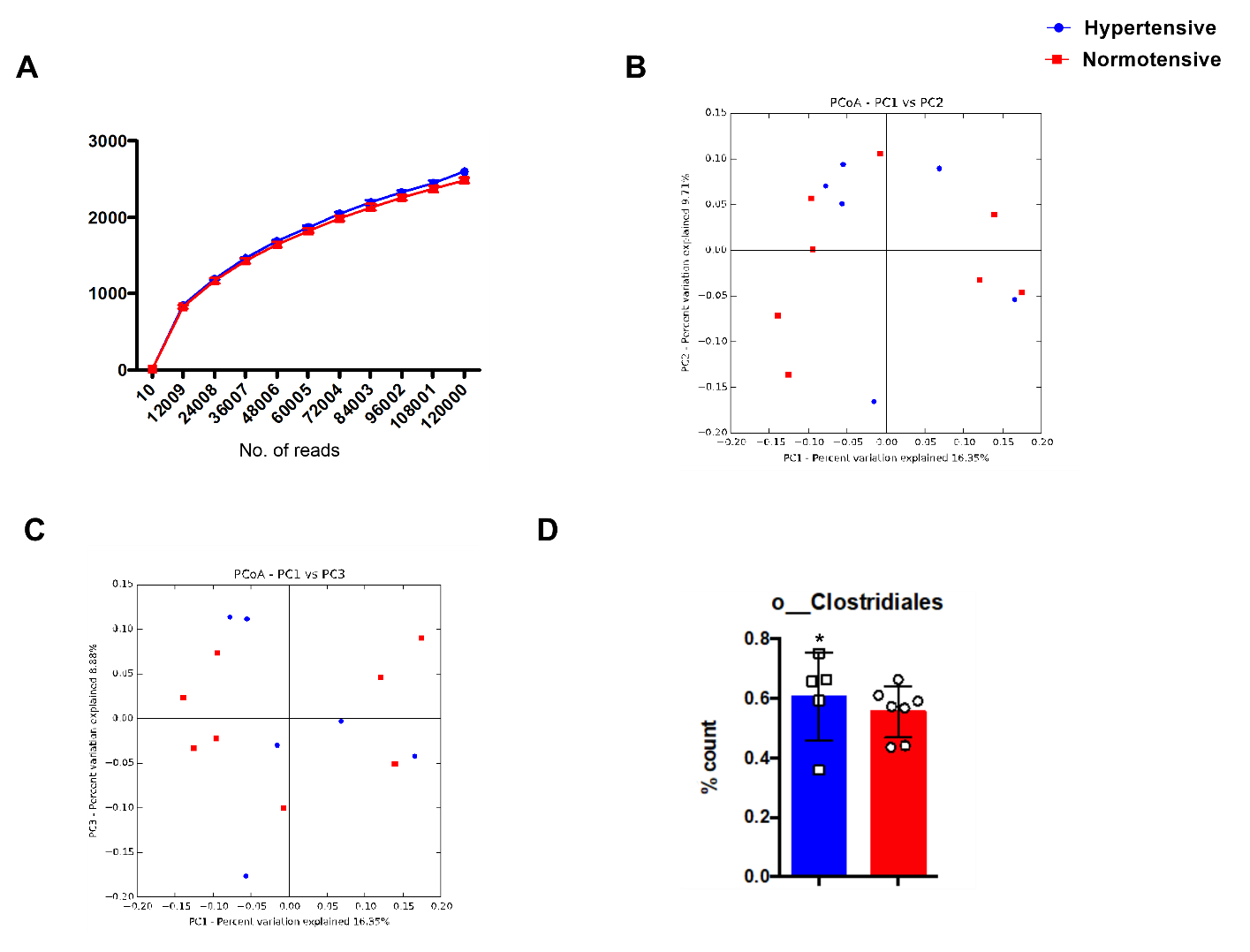

**Figure S3:** Fecal microbiome analysis from the longitudinal study (study 2) showing (**A**) alpha diversity, unweighted beta diversity profiles by principal coordinate analysis (PCoA) with (**B**) PC1 versus PC2 and (**C**) PC1 versus PC3 and (**D)** increases in percent counts of members of opportunistic bacterial order Clostridales between hypertensive and normotensive groups. Values are shown as mean ± SEM. ***** p<0.05.

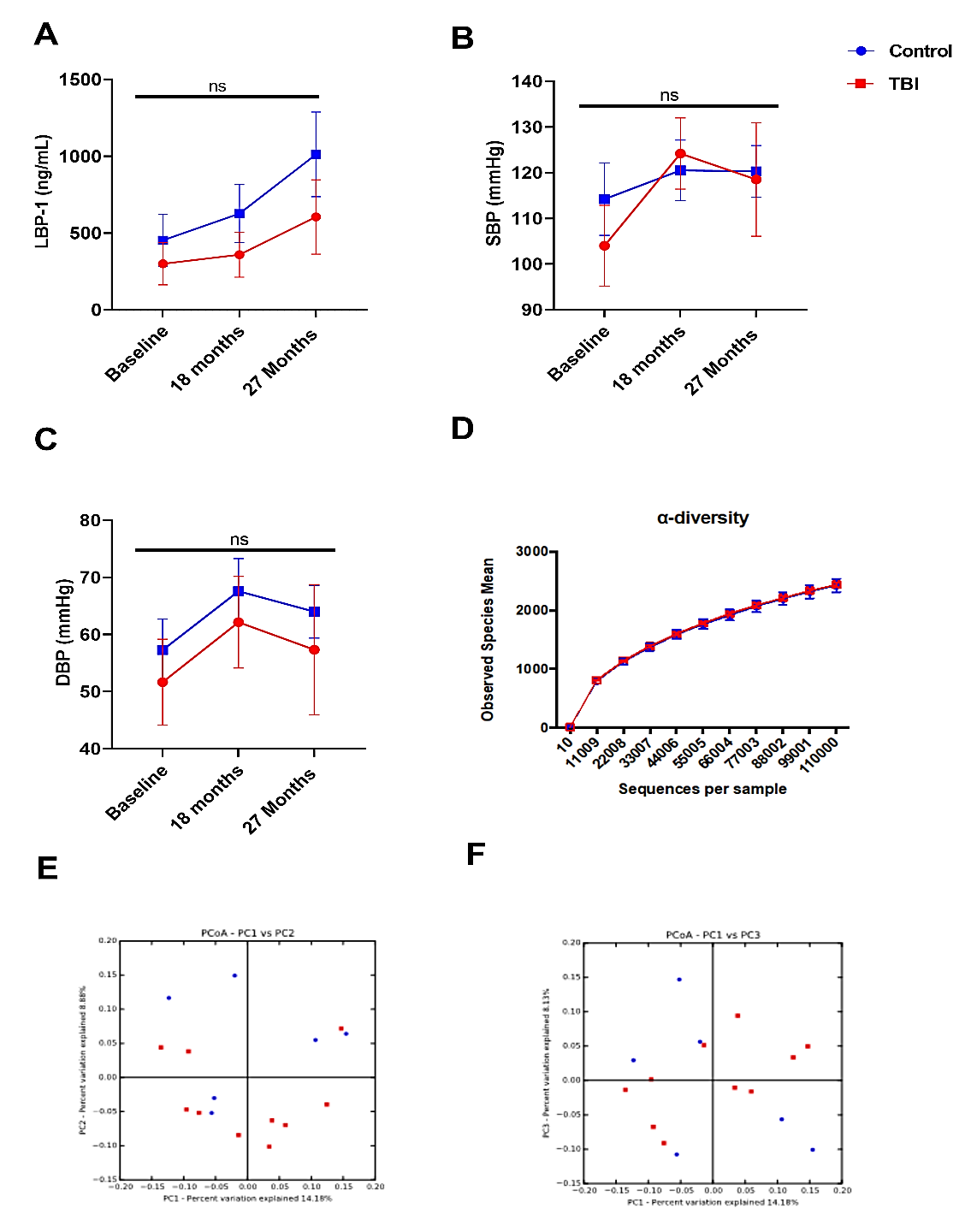

**Figure S4:** No significant differences in (**A**) microbial translocation (LBP), (**B**) systolic blood pressure, (**C**) diastolic blood pressure and fecal microbiome of irradiated cohort from the longitudinal study (study 2) showing (**D**) alpha diversity, unweighted beta diversity profiles by principal coordinate analysis (PCoA) with (**E**) PC1 versus PC2 and (**F**) PC1 versus PC3 no difference between non-total body irradiated controls (non-TBI) and total body irradiated (TBI) animals baseline, 12 months and end of study (27 months). LBP= lipolysaccharide binding protein, SBP= systolic blood pressure, DBP= diastolic blood pressure. ns = non-significant. Values are shown as mean ± SEM.
